# scClassifR: Framework to accurately classify cell types in single-cell RNA-sequencing data

**DOI:** 10.1101/2020.12.22.424025

**Authors:** Vy Nguyen, Johannes Griss

## Abstract

**Motivation:** Automatic cell type identification in scRNA-seq datasets is an essential method to alleviate a key bottleneck in scRNA-seq data analysis. While most existing tools show good sensitivity and specificity in classifying cell types, they often fail to adequately not-classify cells that are not present in the used reference.

**Results:** scClassifR is a novel R package that provides a complete framework to automatically classify cells in scRNA-seq datasets. It supports both Seurat and Bioconductor’s SingleCellExperiment and is thereby compatible with the vast majority of R-based analysis workflows. scClassifR uses hierarchically organised SVMs to distinguish a specific cell type versus all others. It shows comparable or even superior sensitivity and specificity compared to existing tools while being robust in not-classifying unknown cell types. As a unique feature, it reports ambiguous cell assignments, including the respective probabilities. Finally, scClassifR provides dedicated functions to train and evaluate classifiers for additional cell types.

**Availability and Implementation:** scClassifR is freely available on GitHub (https://github.com/grisslab/scClassifR).

## Introduction

Single-cell RNA-sequencing has become a key tool for biomedical research. One of the key steps in analyzing single-cell RNA-sequencing data is to classify the observed cell types.

The most common approach to annotate cell types is using cell clustering and canonical cell type-specific marker genes. However, this has several major drawbacks. First, the work requires profound knowledge on a wide range of cell populations. The situation becomes more complicated if a dataset contains highly similar cell types such as T cells, ILC, and NK cells. Second, cell clusters may not be “pure” but contain mixtures of multiple cell types. Such cases are often missed when only focusing on cluster-specific marker genes. Finally, this manual approach does not efficiently scale to large-scale studies or data reanalysis and is inherently hard to reproduce. Therefore, automated methods are needed to identify cell types in scRNA-seq data.

In recent years, several computational methods were developed to automate cell identification. This includes methods that identify cell types by projecting cells to cell type landmarks, then inferring unknown cells close to already known cell types in the embedded space (northstar [1], scmap [2], MARS [3]). A further approach is to correlate gene expression in annotated groups/clusters of cells with unannotated populations (scCATCH [4], SingleR [5], CIPR [6], clustifyr [7], scMatch [8]). Without using annotated datasets, DigitalCellSorter [9] classifies cells based on the expression of high impact biomarkers, where the impact of the biomarkers depends on their unicity to particular cell types. A large number of algorithms use machine learning (CellAssign [10], SciBet [11], Garnett [12], CHETAH [13], SCINA [14], scPred [15], scID [16]), or neural networks (ACTINN [17], MARS [3]) to automatically learn mapping functions from gene expression of annotated cells to classes of those cells. Despite this large number of cell classification approaches, several approaches show weaknesses that prevent their easy implementation into existing workflows.

We classified existing tools based on key features for efficient automatic cell type classification (Table 1). Only tools that classify individual cells instead of whole clusters can be used to identify issues in the cell clustering. While the vast majority of tools support the identification of new or user supplied cell types (De novo cell type discovery) a considerable portion does not report reliability scores. Moreover, we only identified three tools that are able to report ambiguous cell type assignments, MARS, DigitalCellSorter, and CHETAH. This is crucial since many cell types are closely related, such as monocytes, macrophages, and dendritic cells, which can easily lead to incorrect classification results. Finally, several tools are unable to not-classify cells that are not present in the used reference (unknown population detection). Therefore, we only identified one R based tool that contains all features which we feel are necessary for accurate cell type classification.

**Table 1:**
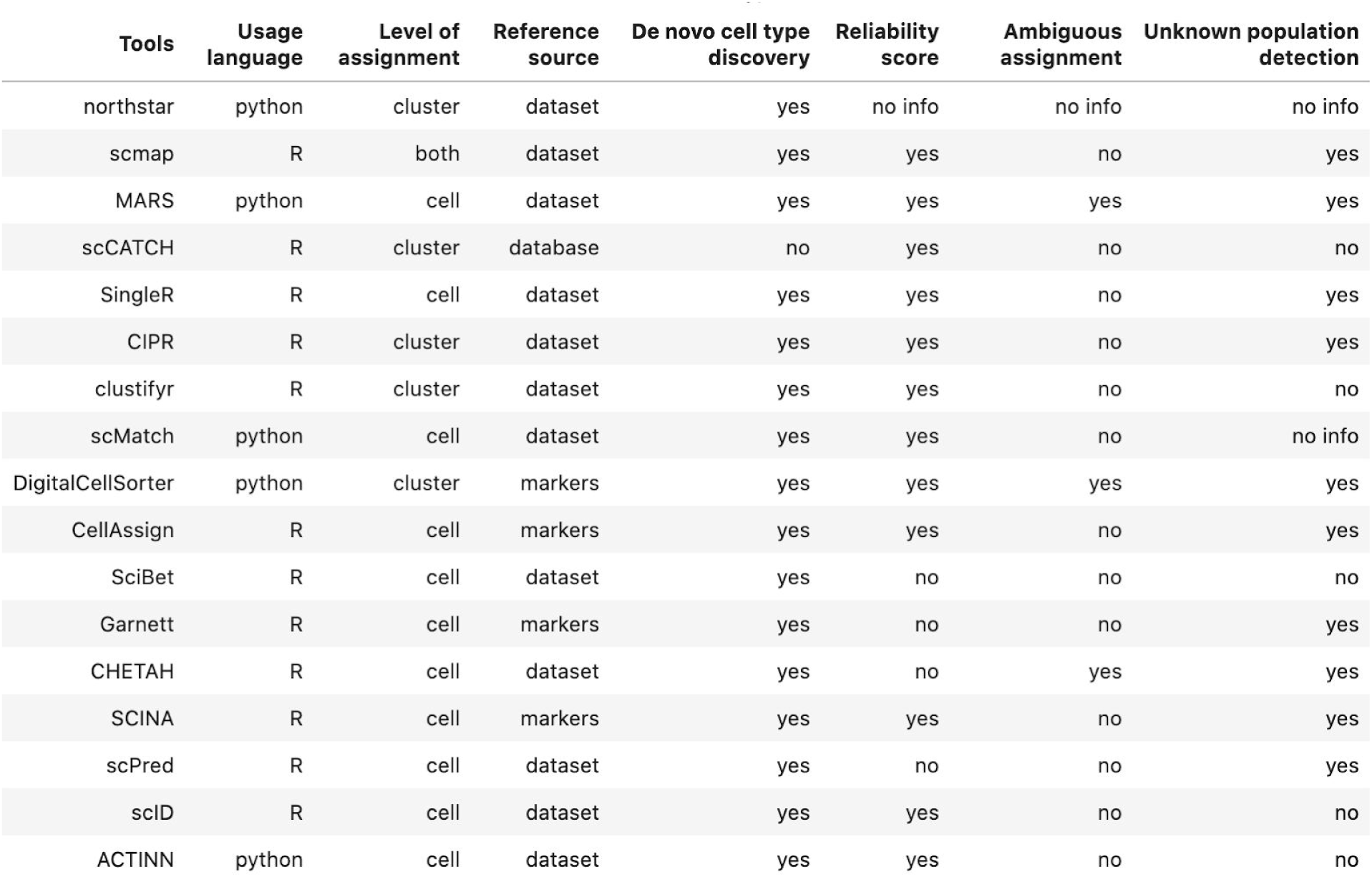
Structured list of existing tools to automatically classify cell types in scRNA-seq datasets.

Here we present scClassifR, a novel tool to automatically classify cells in single-cell RNA-sequencing datasets. scClassifR ships with predefined models for several cell types that can be easily extended by the user.The package uses SVM learning models organised in a tree-like structure to improve the classification of closely related cell types. Most importantly, scClassifR reports classification probabilities for every cell type and reports ambiguous classification results. Therefore, scClassifR fills an important need in the automatic classification of cell types in single-cell RNA-sequencing experiments.

## Implementation

scClassifR is an R Bioconductor package to classify cell types using pre-trained classifiers in scRNA-seq datasets. The package revolves around an S4 class called scClassifR. Each object of the class defines a classifier of a cell type wrapping 5 pieces of information: the classified cell type corresponding to the name of the classifier, a RBF SVM model learned and returned by the caret package [18], a feature set on which the model was trained, a prediction probability threshold and the parent of the classified cell type (if available). Trained models are stored in a named list which are referred to as a classifier database. The package has already a built-in database of pre-trained classifiers which can easily be extended or even replaced by the user.

### Classification process

The core classification task is performed using RBF SVM-based machine learning models on predefined sets of features through the caret R package. C was constantly set at 1, and sigma was defined by the formula below:

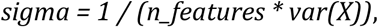

while X was the expression matrix and n_features the number of features used.

scClassifR further supports the concept of “child” and “parent” classifiers. Child classifiers are used to further sub-classify cells that are already classified by a parent classifier. Child classifiers are trained and applied only to cells that were already classified by the respective parent classifier. Internally, scClassifR continuously ensures that the classifier database is consistent and that child classifiers are only added to a database if the respective parent classifier is present. This structure can be visualized using the “visualize_tree” function. Internally, it uses the data.tree R package to plot the hierarchical structure of cell types. Taking the database location provided by users, the package automatically consumes the database, gets classifiers’ information, forms cell type relationships and creates a tree of cell types. The tree has nodes corresponding to main or parent cell types and leaves corresponding to cell types having no children. Thereby, the user can get a quick overview over all available cell types.

scClassifR supports both Seurat and Bioconductor’s SingleCellExperiment class objects as input to its central “classify_cells” function. The classification result is returned as new metadata slots in the input object storing the cell type(s) and the respective classification probabilities. Thereby, the classification results can be directly visualized and further analysed using the respective standard plotting functions and workflows.

### Training new classifiers

scClassifR simplifies the task of training and evaluating new classifiers. The training process (“train_classifier” function) supports both Seurat and SingleCellExperiment objects as input and produces a scClassifR object. The user must specify which features are used for the training process. Generally, we recommend using around 20-30 known canonical marker genes. Increasing this number quickly results in an overfitting of the training data. If non-normalised data is supplied, the train_classifier function can automatically perform a z-score transformation on the input data. A balancing process ensures that an equal number of cells are present in the target and non-target class. Finally, the caret training function is used to train the classifier. Thereby, all steps required to train a new classifier are available in a single, simple to use function.

The testing process through the “test_classifier” function follows a similar process as the training function. Using an independent test dataset as input, the test_classifer function calculates an overall AUC score, the accuracy, sensitivity and specificity of the classifier at the defined probability threshold and at thresholds from 0.1 to 0.9 with steps of 0.1 to simplify the tuning of the probability threshold. Once the classification model meets the user’s expectations, scClassifR provides several functions to store the classifiers in a common database. These functions ensure that the database remains consistent with respect to parent and child classifiers. Thereby, scClassifR provides a complete infrastructure to train and evaluate new classifiers.

## Methods

### Datasets

We used twelve public scRNA-seq datasets for the creation of the in-built classifiers and the benchmarking of the package. These are two melanoma datasets from Sade-Feldman *et al.* (GSE120575)[19] and Jerby-Arnon *et al.* (GSE115978)[20], and five pancreas datasets from Baron *et al.* (GSE84133)[21], Muraro *et al.* (GSE85241)[22], Segerstolpe *et al.* (E-MTAB-5061) [23], Wang *et al.* (GSE83139)[24] and Xin *et al.* (GSE81608)[25].

Additionally, five datasets were used to evaluate the performance on closely related cell types: PBMC 3k [26], PBMC 500[27], PBMC - Ding SM2 [28][29], HIV1 [30], and Lung - Zilionis [31][32].

### Data preprocessing and cell type assignment

scRNA-seq data was downloaded from GEO for the Sade-Feldman and the Jerby-Arnon datasets (TPM counts). For the Sade-Feldman melanoma dataset, we preprocessed the data following the authors’ approach: First, we filtered out mitochondrial genes. We then retrieved only cells expressing at least 1000 features and only features expressed in at least 3 cells. Finally, we kept only cells with housekeeping genes expressed at low levels: log2(TPM + 1) < 2.5. For the Jerby-Arnon dataset, we eliminated mitochondrial genes, and filtered out cells having less than 1000 expressed genes and genes expressing in less than 3 cells. The datasets were then normalized and scaled using the basic pipeline of Seurat v3 regressing out confounders, such as patients (for Sade-Feldman dataset), samples and cohorts (for Jerby-Arnon dataset). After that, the data dimension was reduced to the first 40 (Sade-Feldman dataset) and 45 (Jerby-Arnon dataset) principal components. lustering was performed with default parameters.

The five pancreas scRNA-seq datasets were preprocessed based on the protocol proposed by the Hemberg lab [33]. The datasets were then normalized and scaled following v3 Seurat SCTransform protocol with regressing out main confounders: samples, patients/donors, diseases/conditions, and batches. Number of principal components for the five datasets (Baron, Muraro, Segerstolpe, Wang and Xin) are 45, 40, 45, 40, and 30, respectively. Nearest neighbors and clusters were computed using the default parameters, except clusters in Wang dataset were calculated at resolution = 1.

Cells in Baron and Wang datasets were annotated by authors. Cell types in three other datasets were manually assigned on the cluster-level and based on known canonical markers (Supplementary Table 1).

The PBMC 3k was analyzed using Seurat v3.1 following the respective vignette [26]. The PBMC 500 dataset was preprocessed and analyzed according to the ILoReg v1.0 vignette [27]. The PBMC - Ding SM2 was downloaded from the Single Cell Portal [29]. Here, we only retained the data sequenced using the Smart-seq2 protocol. This subset was then preprocessed using Seurat’s basic analysis pipeline following the SCTransform strategy and regressing out the experiment property. After that, 20 principal components were used for clustering. Original cell labels from the authors were used.

The HIV1 dataset was downloaded from the Single Cell Portal and processed using Seurat’s scTransform pipeline regressing out the donor identification [34]. 50 principal components were used for clustering and the cell labels were provided by authors and integrated into our Seurat object.

The human lung dataset by Zilionis *et al*. [31] is available through the scRNAseq R package [32]. The SingleCellExperiment object was converted to a Seurat object and processed using the Seurat analysis pipeline with SCTransform. Patient and tissue features were regressed out. 50 principal components participated in the neighbors and clusters finding process. Cell labels provided by authors were moved from the SingleCellExperiment metadata to the Seurat metadata slot.

### Pretrained learning models for basic immune cells

We used the Sade-Feldman scRNA-seq dataset for training and the Jerby-Arnon dataset to test our package’s inbuilt classifiers. Training and testing of classifiers was performed using the package’s inbuilt functions (see above). At the time of writing, the package contains classifiers for B cells, plasma cells, NK, T cells, CD4+ T cells, CD8+ T cells, monocytes, dendritic cells, and pancreatic alpha, beta, gamma, delta, ductal, and acinar cells.

### Benchmarking discrete populations using the pancreas datasets

This benchmark was performed using the 5 pancreas datasets and a 5-fold cross-validation scheme. In each fold, one among five datasets was used for training and the other four for testing.

Preprocessed and analyzed Seurat objects were converted into SingleCellExperiment objects (for clustifyr and SingleR), into expression matrices (for SciBet) or into CellDataSet objects (for garnett).

For Garnett and scClassifR, we used the same set of features for training selecting features fo six cell types: alpha, beta, delta, gamma, ductal, and acinar, were prepared. For SCINA we use Seurat’s FindAllMarkers function to get the top 10 differentially expressed markers of the respective clusters in the training dataset. These were then used as a signature. For all other methods we used the standard pipeline provided by the authors for training and testing.

The benchmarking process assessed two aspects of the cell classification accuracy: 1) How well a specific cell type was recognized and 2) how often cell types were misclassified that were not part of the reference.

The accuracy of specific cell type identifications was assessed by transforming the classification results into a binary matrix where each cell type is represented as one column and each cell as one row. For each cell we record whether it was classified as that specific cell type (1) or not (0). As we know the correct number of specific cells in the training dataset, we can then calculate the sensitivity and specificity per cell type. Summary statistics are then reported as the average sensitivity and specificity across all classified cell types. scClassifR can report ambiguous cell types. In order to ensure a fair comparison between the different tools, only the most probable cell type (as implemented in the scClassifR package) was used for the benchmark.

The misclassification of cell types was assessed by testing how cell types that are not part of the training dataset were classified. This rate was defined as the ratio of cells that remained unclassified divided by the total number of cells that should not have been classified. Here, we refer to this rate as the ‘unknown population detection rate’.

### Benchmarking closely related populations

This benchmark was performed on five datasets: the PBMC 3k dataset analyzed by Seurat v3.1 [26]; the PBMC 500 dataset, which was a subset of the PBMC 3k dataset but preprocessed and analyzed by ILoReg v1.0 [27]; the PBMC dataset by Ding *et al*. [28], the subset sequenced by Smart-seq2; the hyper acute HIV1 dataset by Kazer *et al*. [30], available on the Single Cell Portal [34]; and the human lung dataset by Zilionis *et al*. [31], available in the scRNAseq R package v2.4 [32].

scClassifR used in-built classifiers for B cells, T cells, NK, which were trained on the Sade-Feldman melanoma dataset and tested on the Jerby-Arnon melanoma dataset. SingleR, clustifyr, SciBet and Garnett were trained on the Sade-Feldman melanoma dataset. Garnett and SCINA require a set of markers as input. Therefore, we choose two sets of markers for them: the set of markers used by our in-built classifiers (Garnett-1 and SCINA-1 as results), and the biomarkers in the Sade-Feldman dataset analyzed by the FindAllMarkers function of Seurat v3.1.

Cell types were classified in each dataset and the sensitivity and specificity calculated per cell type as described above. Results are reported as average across all cell types per dataset. For each dataset, we further assessed the unknown population detection rate (see above).

### Runtime estimation

For both benchmarks we recorded the runtime of the classification tools as wall clock time. The exact moment of start and end were retrieved using the sys.time() function in the R base package.

## Results

scClassifR is compatible with both Seurat and Bioconductor’s SingleCellExperiment object. and ships with pre-trained classification models for most basic immune cells. Therefore, it can easily be integrated into the vast majority of existing scRNA-seq workflows.

scClassifR uses SVMs to classify cells. These are wrapped by the new R S4 object “scClassifR” together with all required parameters to apply the model. Classifiers are stored in a hierarchical tree-based structure allowing the definition of “parent” and “child” classifiers. In such cases, cells are first classified using the parent classifier. Only cells identified as that specific cell type are then further classified using the respective child classifier. The complete collection of models in the tree based structure is stored in a single file.

Cell classification results are directly stored in the metadata slots of the respective SingleCellExperiment or Seurat object. This includes possible ambiguous cell type assignments, as well as the classification probability for every cell type. Thereby, all results produced by scClassifR can be immediately visualized and further analysed using the respective pipelines default functions.

Finally, scClassifR offers a user-friendly environment to train and test new cell classifiers (see Methods for details). All functional parameters are adjustable and configurable, which gives the user full control during the training process. Thereby, scClassifR offers a complete framework for the automatic classification of cell types in scRNA-seq datasets.

### Hierarchical classification models help identify unrecognized sub-populations

A key challenge in the characterisation of cell types in scRNA-seq datasets is to what level of detail cell types should be classified. Several research questions focus on very specific subtypes, for example specific B cell phenotypes. At the same time, other B cell subtypes may be of less interest - or be unexpected at all. In tools that do not support hierarchical classification models researchers have to either classify all B cells at the same level of detail (with the danger of missing rare subtypes) or leave a large portion of cells unclassified.

scClassifR’s hierarchical organisation of cell classification models is ideally suited for such targeted cell classification approaches. First, researchers can train a parent classifier to identify all cells belonging to the general cell type of interest. In a second step, they can now create a child classifier(s) to focus on their subtype(s) of interest. Figure 1 highlights these two approaches. scClassifR’s inbuilt classifiers contain a hierarchical model for overall “B cells” and its child terminally differentiated “plasma cells”. Figure 1A highlights that the dataset contains several plasma cells, but a large portion of the general “B cells” is only captured by the parent classifier. If we were to focus purely on plasma cells and only train a respective plasma cell classifier, a large portion of B cells remained unclassified which may lead to a misinterpretation of the results (Figure 1B). Additionally, a group of cancer-associated fibroblasts (lower left group of cells, Figure 1B) were misclassified as plasma cells. These express SDC1, a sensitive but not specific plasma cell marker. Due to their additional expression of FAP, PDGFRA, PDGFRB, TAGLN, and COL1A1 we can be certain that they are not plasma cells. The general B cell classifier was able to correctly distinguish these cells (Figure 1A). This example highlights that scClassifR’s hierarchical classification system is ideally suited to classify cells at a high level of detail.

**Figure 1:**
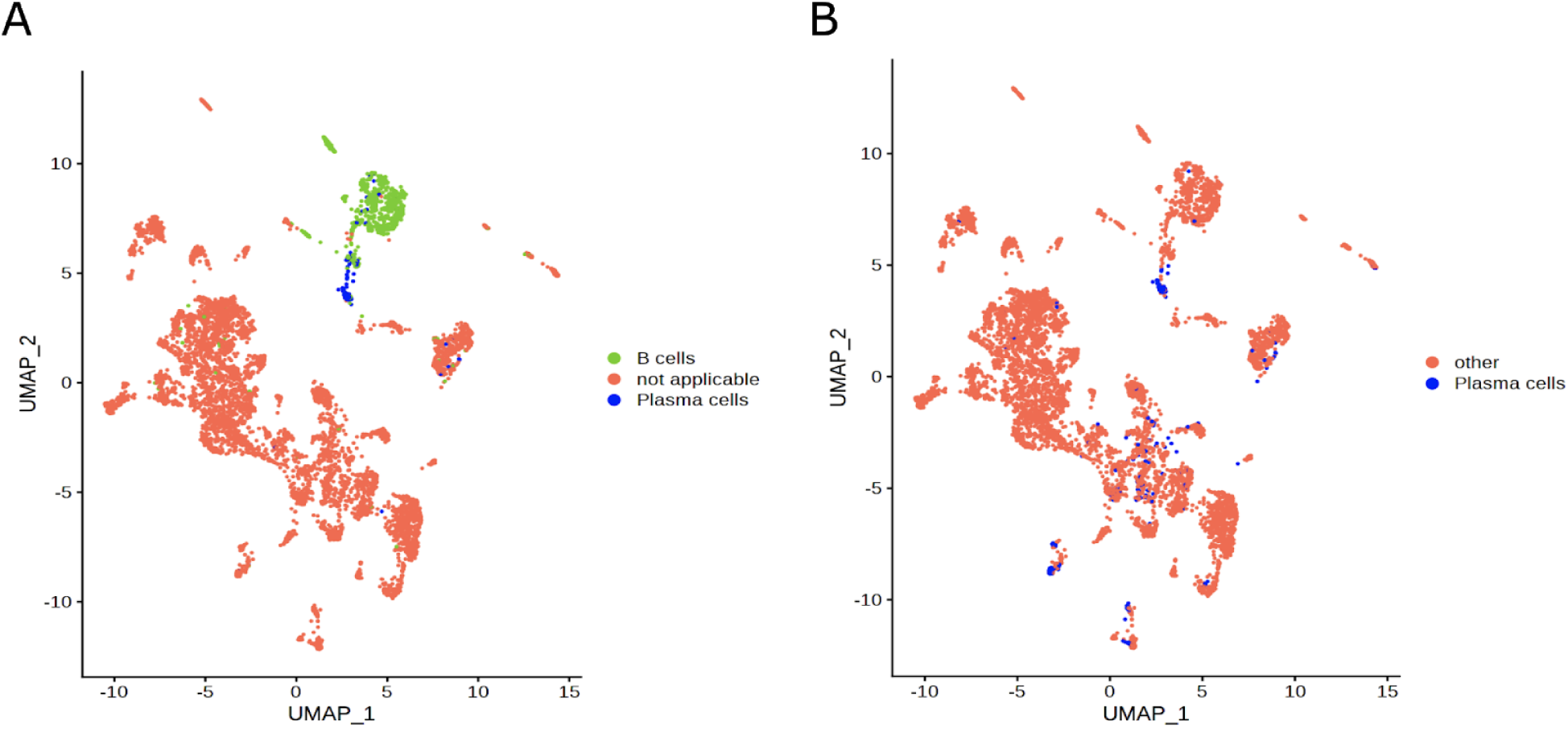
UMAP plot displaying the classification results in the Jerby-Arnon melanoma dataset. **A)** Classification results of a plasma cell classifier as a child of the more general B cell classifier. **B)** Classification results of a plasma cell classifier trained separately without any parent classifier.

### Prediction by scClassifR is compatible with Seurat clustering and biomarkers

First, we compared scClassifR’s automatic cell identification with manual cell type assignments based on cell clustering results and expression of canonical markers. Here, we used the Muraro *et al.* pancreas dataset [22]. First, we analysed the dataset through the Seurat v3.1 analysis pipeline. We then classified the resulting cell clusters based on known canonical markers (Supplementary Table 1) which was in-line with the authors original assignment [22]. Next, we trained 6 new models classifying alpha, beta, delta, gamma, ductal and acinar cells on the Baron *et al.* dataset [21]. The first 4 models were tested on the Xin *et al.* dataset [25] and the last two on the Wang *et al.* dataset [24]. scClassifR’s classification derived from completely independent datasets was perfectly in-line with the manually derived ones (Figure 2). Cell types not present in the training dataset (endothelial and mesenchymal cells) were further correctly not classified and labelled as “unknown” cells. This highlights that scClassifR can robustly classify cell types in independent datasets.

**Figure 2:**
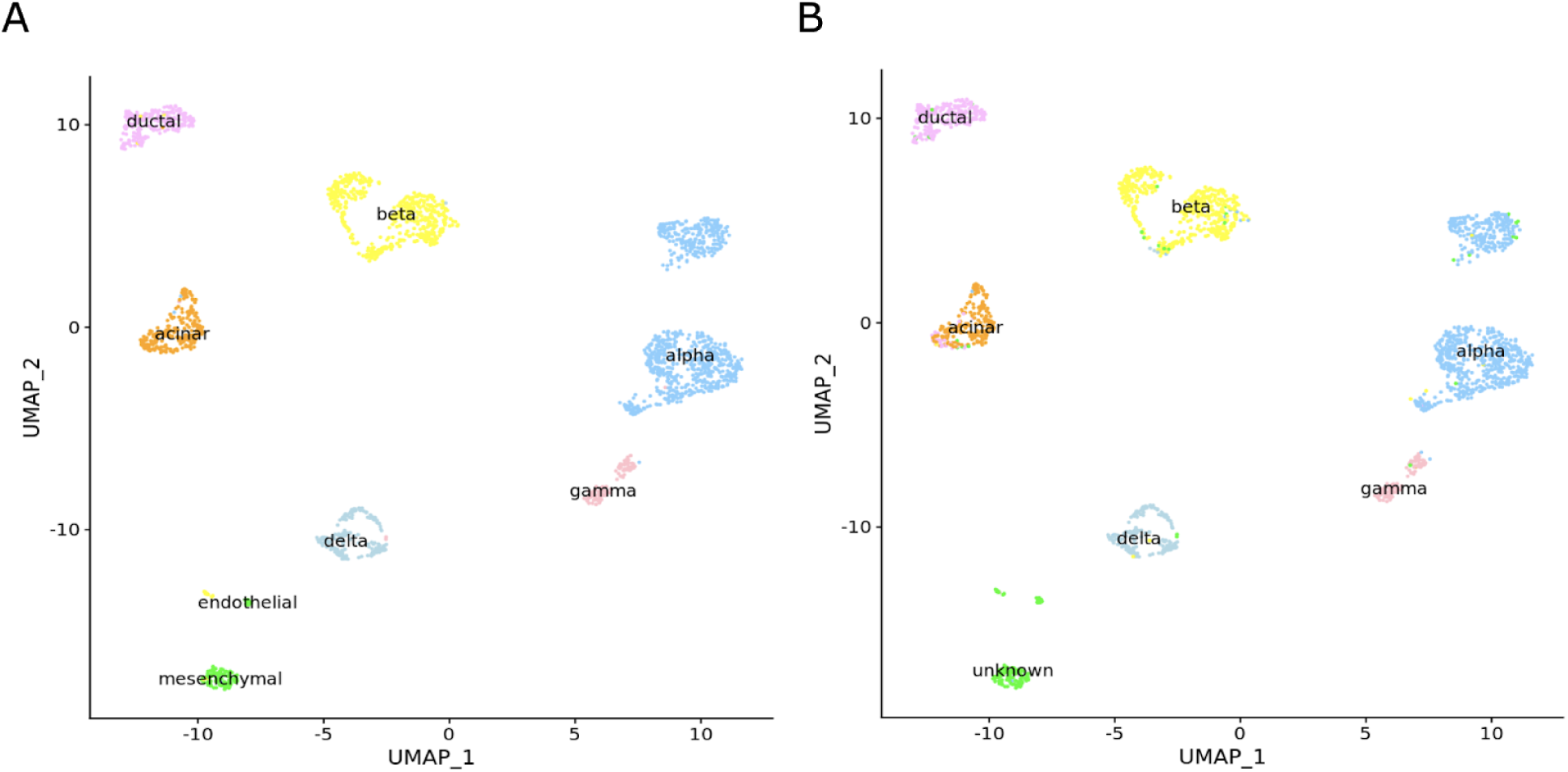
Classification of cell types in the pancreas dataset from Muraro *et al.* based on canonical markers and manual assignment (**A**) and an automatic classification using scClassifR (**B**) where cell types were learned from the independent Baron *et al.* dataset. Cell types not present in the training data (mesenchymal and endothelial cells) were accurately classified as “unknown” cell types.

### scClassifR can reveal inaccurate clustering results

Since scClassifR classifies individual cells, the classification results can be used to cross-validate the clustering results. We therefore performed an additional benchmark to evaluate scClassifR’s ability to determine cell types in such mixed clusters (Figure 3). A CD8+ T cell model was trained as a child model of T cells on the Sade-Feldman melanoma dataset and tested on the Jerby-Arnon melanoma dataset. The cell labels in the Jerby-Arnon datasets were previously assigned based on the clustering results (Figure 3B). This results in an AUC score of 0.85, which is acceptable but not ideal. A more detailed look at the expression of canonical CD8+ T cell markers reveals that a subset of cells does not express CD8A and CD8B (Figure 3A). These cells were all merged into a single cluster by the default Seurat pipeline and the cluster could not be attributed to a specific T cell phenotype due to the inconsistent marker expression (Figure 3B). scClassifR’s classification results accurately reflected this variation in marker expression (Figure 3C). Thereby, scClassifR is able to highlight even fine inaccuracies of the cell clustering results.

**Figure 3:**
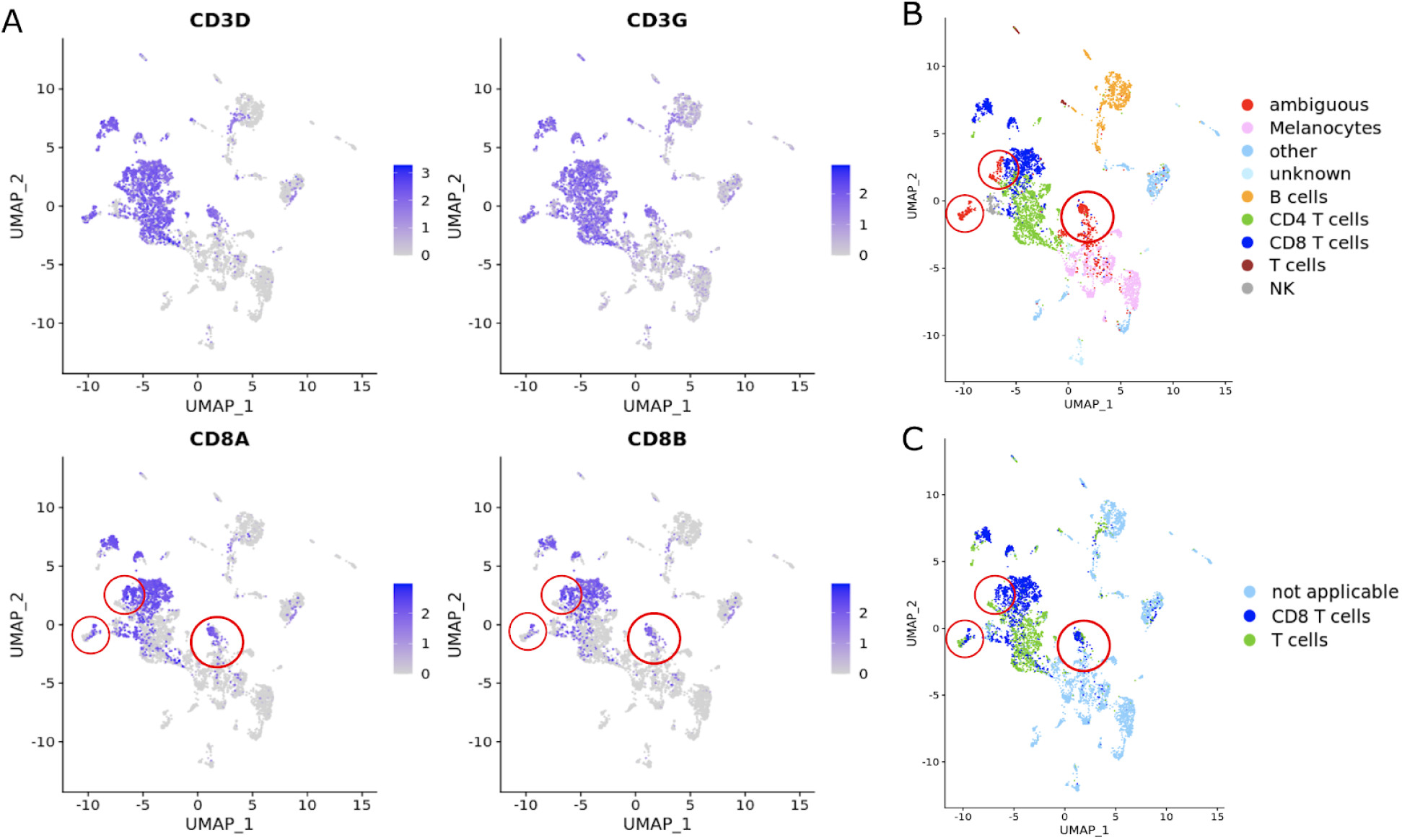
UMAP plot of the Jerby-Arnon dataset showing specific T cell marker expression (**B**) and the manual, clustering based (**B**) and automatic scClassifR based (**C**) classification of cell types. Red circles highlight areas where the clustering results merged T cells with other cell types.

### scClassifR outperforms existing tools considering sensitivity, specificity, and the detection of unknown cell types

An important aspect of automatic cell type classification tools is the ability to correctly deal with unknown cells. Reference datasets may always be incomplete. Therefore, tools need to be able to recognize such unknown cells to avoid a misinterpretation of the data.

We compared scClassifR’s performance against existing tools using two benchmarks: first, a group of datasets containing very discrete cell populations and second a group with closely related immune cell populations. The benchmarks included six existing tools. SingleR [5] selects the most variable genes for each cell type in an annotated dataset. Then, cell types are identified in an unlabelled dataset by correlating the expression values. CHETAH [13] selects the top differentially expressed genes and finds the distribution of correlation between cells in each cell type, unknown cells are then classified by the high cumulative density of a cell type correlation distribution. SciBet [11] retrieves cell type markers and eliminates noisy genes by the E-test. For each cell type, SciBet learns a multinomial model to form a likelihood function defining the probability of each cell to belong to a cell type, hence cell annotation relies on a likelihood maximization process. Without having a marker identification process, Garnett [12] requires a list of marker genes as input to train cell classifiers. clustifyr [7] is the only tool working on the cluster level. It identifies cell types through the correlation of cluster gene expression with annotated cell expression values. SCINA [14] relies on user-supplied marker genes to assign cell types and depends on the users’ prior knowledge. Thereby, we get a comprehensive assessment of scClassifR’s performance in comparison to existing tools.

#### Classifying discrete cell populations

We performed the benchmark using a five-fold cross-validation scheme with five pancreas datasets [21–25]. In each fold, one of the five datasets was used for training, the other four for testing. In each iteration, we assessed the sensitivity and specificity for each classified cell type (see Methods). Results are reported as the average sensitivity, and specificity across all cell types for each iteration (Fig. 4A, B). Additionally, we assessed the unknown population detection rate, which is defined as the number of correctly unassigned cells over the total number of cells that are not present in the reference (Fig. 4C, Supplementary Table 2). scClassifR and CHETAH are the only evaluated tools able to return ambiguous cell type assignments. In order to ensure a fair comparison, only the most probable cell type reported by scClassifR was used for the benchmark. As CHETAH does not provide a reliability score for ambiguous cells, these had to be excluded for the benchmark. This five-fold cross-validation benchmark thereby ensures that we get an accurate and comparable estimate of each tool’s performance.

**Figure 4:**
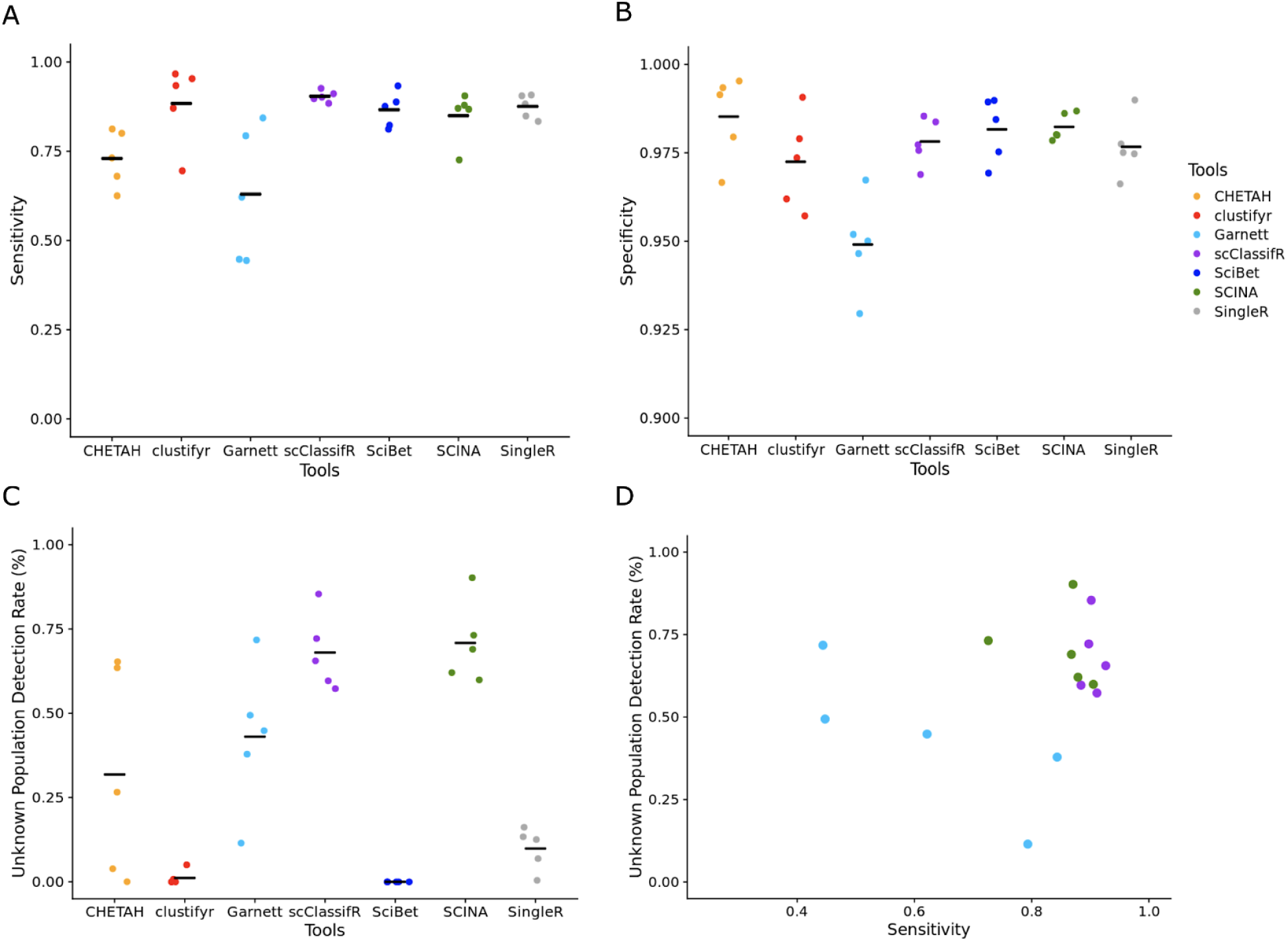
Benchmark results when classifying distinct cell populations in five pancreas datasets. The benchmark was performed in a five-fold cross-validation scheme where one dataset was used for training and the other four for testing. The shown values represent the mean across all classified cell types across the four evaluated datasets per iteration. Panels show the sensitivity (**A**), specificity (**B**), and the proportion of cells not present in the training data, that were correctly not-classified (**C**). **D**) The relationship between the proportion of correctly not-classified cells versus the sensitivity for Garnett and scClassifR.

Throughout all iterations, scClassifR was consistently among the tools with the highest sensitivity and specificity, but next to SCINA the only tool with an acceptable unknown population detection rate (Figure 4A-C, Supplementary data 1). CHETAH’s low sensitivity comes from its ambiguous cell type assignment without information about the most probable cell type. Therefore, these cells had to be excluded from the analysis. Including them would have increased CHETAH’s sensitivity but lead to a considerably worse specificity. The fact that clustifyr works on the cluster level leads to a win all or lose all scenario. In datasets where the clustering results were suboptimal, clustifyr’s performance decreases dramatically. Garnett’s increased detection of unknown cell types comes at a direct cost of reduced sensitivity (Figure 4D). This is not the case for scClassifR and SCINA which both showed comparable unknown detection rates with a stable sensitivity. Overall, scClassifR showed the highest sensitivity across all tools while being able to correctly recognize unknown cells.

#### Closely related populations

Our second comparative benchmark focuses on the differentiation of closely related immune cell types. Here we used five annotated datasets: the PBMC 3k dataset as analyzed in the Seurat v3.1 tutorial [26], the PBMC 500 dataset analyzed by ILoReg v1.0 tutorial [27], the SM2 PBMC dataset as part of the PBMC dataset by Ding *et al*. [28][29], the hyper acute HIV1 dataset [30][34], and the human lung dataset by Zilionis *et al.* [31] in the scRNAseq package v2.4 [32]. The selection of datasets for this benchmark ensures that we can assess the classification performance in closely related cell types.

Training data was selected to ensure a fair comparison between all tools. scClassifR uses the in-built classifiers for T cells, B cells, NK cells, monocytes (macrophages), and dendritic cells, which were trained on the Sade-Feldman melanoma dataset and tested on the Jerby-Arnon melanoma dataset (except dendritic cells which were tested on Butler *et al.* PBMC dataset [35]). SingleR, clustifyr, Garnett and SciBet also used the Sade-Feldman dataset for training. For Garnett and SCINA we evaluated two settings: using the same markers as our in-built classifiers (Garnett-1 and SCINA-1), and using the cell type markers found through Seurat’s findClusterMarker function in the Sade-Feldman dataset (Garnett-2 and SCINA-2). Thereby, we arrive at a fair comparison between all tools in classifying closely related cell types.

scClassifR again showed the highest average sensitivity in this benchmark (Figure 5A). SCINA (1 and 2) was the only tool with a larger variance in sensitivity across the tested datasets. SingleR, clustifyr and SciBet were unable to distinguish monocytes versus dendritic cells and classified almost all dendritic cells as monocytes (Supplementary data 2). Garnett-1 and SCINA-1, which used the same markers as scClassifR’s in-built classifiers, identified more cells correctly than Garnett-2 and SCINA-2, which used the differentially expressed genes from the Sade-Feldman dataset. SCINA (1 and 2) and scClassifR showed the best specificity (Fig. 5B). Therefore, scClassifR was again among the top-performing tools in terms of sensitivity and specificity when classifying closely related cell types.

**Figure 5:**
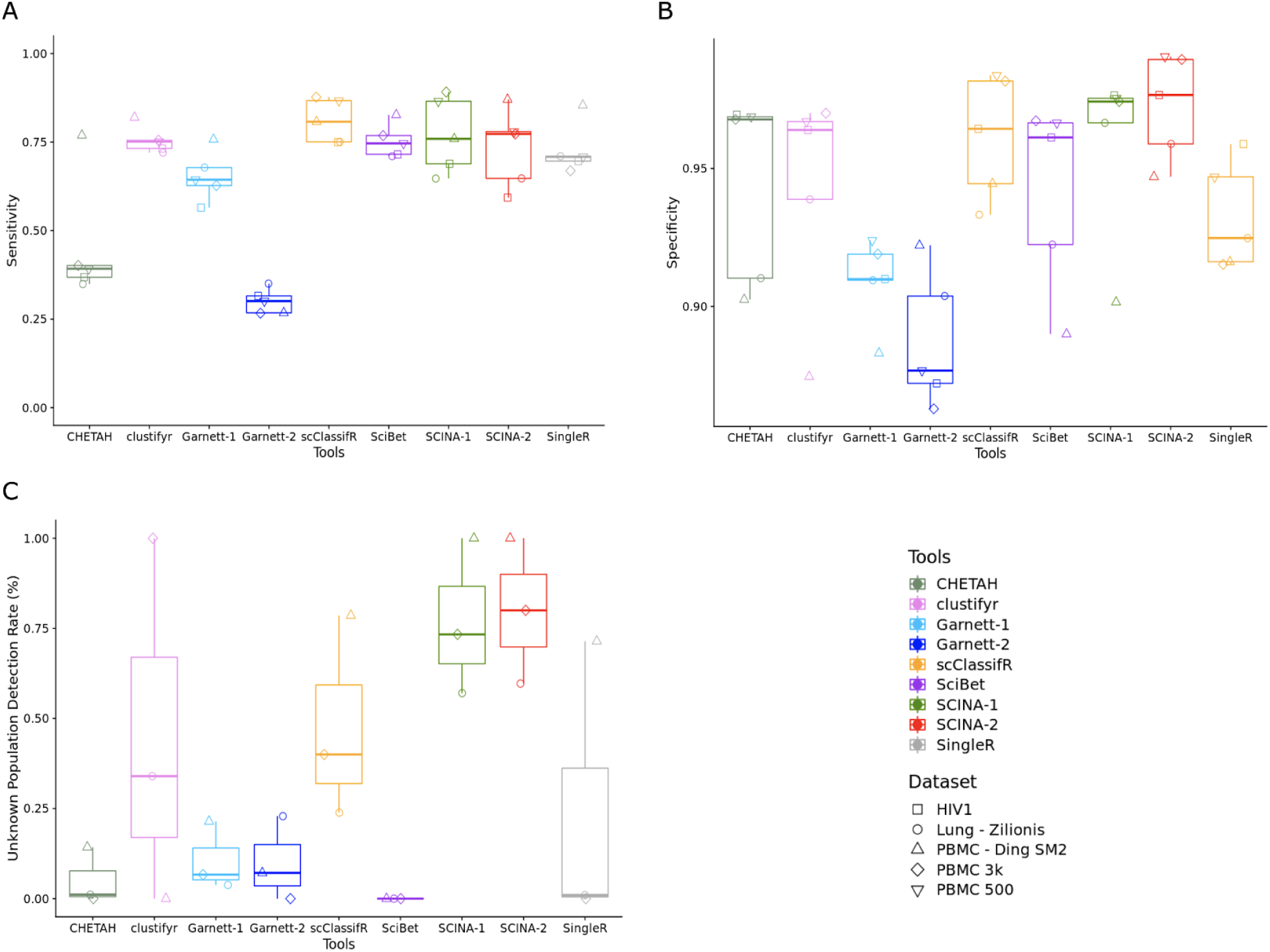
Benchmark evaluating classification accuracy for closely related immune cell types in five datasets. All tools were trained on the independent Sade-Feldmann melanoma dataset. Garnet-1 and SCINA-1 used the same markers as scClassifR for its training process. Garnett-2 and SCINA-2 used the top marker genes identified through Seurat in the Sade-Feldman dataset for the respective cell types. Values represent the mean across all classified cell types for the specific tool and dataset. Panels display the sensitivity (**A**), specificity (**B**), and the proportion of correctly not-identified cells (**C**).

Overall, more tools were able to detect unknown cells compared to the previous benchmark. SCINA (1 and 2) was able to detect most unknown cell types correctly, followed by scClassifR and clustifyr (Fig. 5C). Garnett (1 and 2), SciBet, and SingleR were once more not able to accurately detect unknown cell types. clustifyr’s performance depended on the accuracy of the clustering results which led to no detected unknown cell types in the PBMC - Ding SM2 dataset but very good performance in the PBMC 3k dataset. Altogether, in this benchmark only scClassifR showed acceptable performance across all measured parameters.

### scClassifR is computationally efficient in large datasets

We estimated the runtime of all applications as wall clock time in the two benchmarks (Fig. 6). The evaluated dataset sizes range from 500 to roughly 60,000 cells and were generally smaller in the first benchmark (Figure 6A). Here, we calculated the average runtime across the five-fold iterations. A 120GB RAM Ubuntu 20.04 machine with 32 cores was used to estimate the runtime in all classification. Thereby, we can estimate the overall performance of each tool.

**Figure 6:**
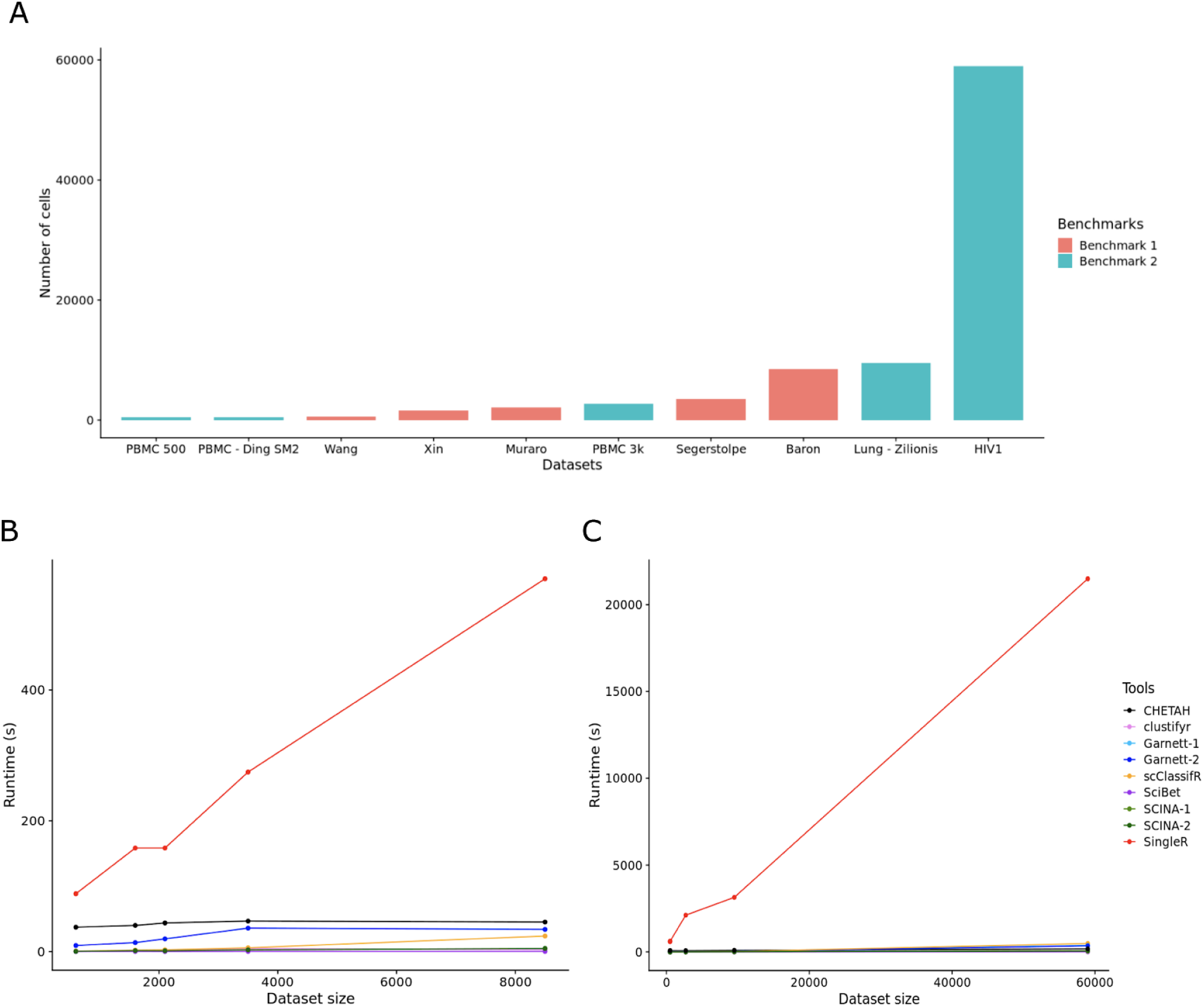
**A)** Size of all evaluated dataset as the number of cells. Runtime as wall clock time for all datasets of the first (**B**) and second (**C**) benchmark.

All tools with the exception of SingleR showed a comparable performance (Figure 6C, D). Only SingleR was significantly slower than the other tools. In the first benchmark, the maximum runtime of SingleR was ~17 minutes when trained on the Segerstolpe dataset and tested on the Baron dataset. Therefore, scClassifR is among the most computationally efficient tools even when analysing large datasets.

## Discussion

Automatic cell type identification in scRNA-seq datasets has become a highly active field and is an essential method to alleviate a key bottleneck in scRNA-seq data analysis. Our benchmarks showed that many of the available tools fail in not-classifying cells that are not present in the training data. This can gravely impact the interpretation of scRNA-seq datasets and results. In our benchmarks, only scClassifR and SCINA were able to consistently achieve a high specificity and sensitivity in both benchmarks while being able to not-classify unknown cell types. scClassifR’s training-based approach led to a more stable high-sensitivity compared to SCINA, which showed the largest variance in our second benchmark. Therefore, scClassifR is a key addition to existing methods to automatically classify cells with a reduced risk to misclassify unknown cells.

A large group of algorithms, such as MARS [3] or SingleR [5], rely on annotated reference datasets. In our experience, this approach is often limited since a single dataset may not contain all cell types of interest. When multiple datasets have to be merged, data size and computationally cost quickly increase dramatically as shown for SingleR in our benchmark. Additionally, sharing annotated reference dataset is complicated by their size. The advantage of scClassifR and other related tools is that the cell type’s properties are learned from a reference dataset, but the reference dataset is no longer necessary to apply the model. This makes the learned models easily transferable, shareable, and reproducible as highlighted by the models shipped as part of scClassifR.

The scarcity of scRNA-seq data often does not permit a clear attribution of a given cell to a specific cell type. When reviewing existing tools, we only found three existing tools that are able to report ambiguous cell type assignments: MARS, DigitalCellSorter, and CHETAH. Ambiguous cell type assignments can quickly allow a researcher to recognise problematic clustering results but can also highlight the limitation in some datasets to clearly identify specific cell types. Therefore, we are convinced that ambiguous cell type identification is a key feature for automatic scRNA-seq cell type assignments.

Finally, scClassifR is among the few tools to provide a dedicated infrastructure to train new cell classifiers. It is impossible to create references that suit all experimental designs. We explicitly provide functions that greatly simplify the training and, most importantly, evaluation of new cell types. Plans are under way to support a GitHub-based central repository for cell type classifiers that also supports multiple species. This will help researchers to quickly share their own classifiers. scClassifR therefore is a scalable, accurate and reproducible method to automatically classify cell types in scRNA-seq datasets.

## Data availability

scClassifR is freely available as open source software on GitHub at https://github.com/grisslab/scClassifR.

## Acknowledgements

This work was supported by FWF-Austrian Science Fund (projects P30325-B28 and P31127-B28).

