## Supplementary Data for "scClassifR: Framework to accurately classify cell types in single-cell RNA-sequencing data"

#### Table of contents

**Supplementary Figure 1** 2

**Supplementary Table 1** 3

**Supplementary Table 2** 4

**Supplementary Table 3** 5

**Supplementary Table 4** 6

### Supplementary Figure 1

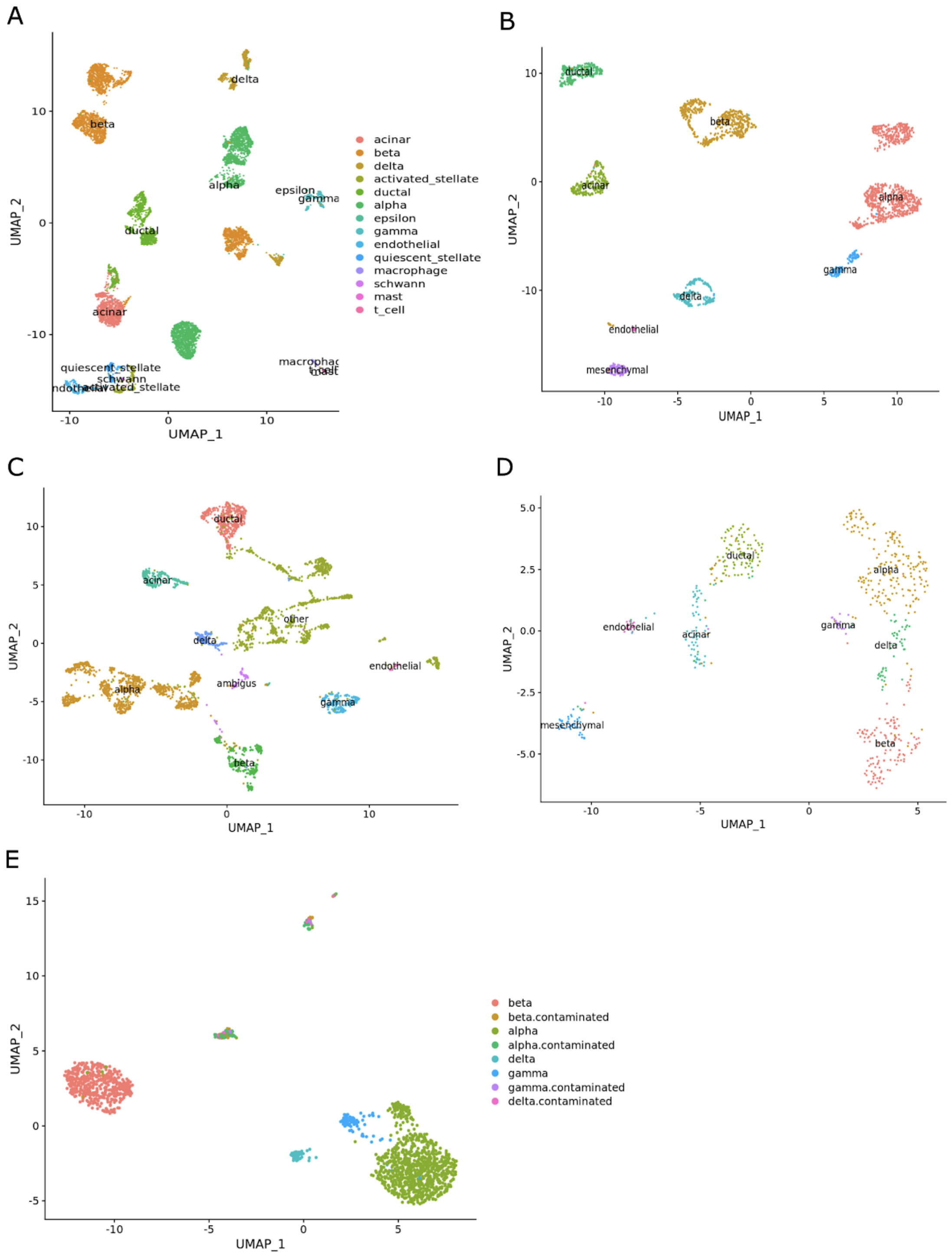


Cell type annotation in pancreas datasets: (**A**) Baron *et al.*, (**B**) Muraro *et al.*, (**C**) Segerstolpe *et al.*, (**D**) Wang *et al.*, (**E**) Xin *et al.*

### Supplementary Table 1

| Cell types | Markers |
|---|---|
| B cells | CD19, MS4A1, CD79A, CD79B, SDC1 |
| Plasma cells | CD19, SDC1 |
| T cells | CD3D, CD3E, CD3G, CD8A, CD8B, CD4 |
| CD4+ T cells | CD3D, CD3E, CD3G, CD4 |
| CD8+ T cells | CD3D, CD3E, CD3G, CD8A, CD8B |
| NK | CD2, NCAM1, NCR1, KLRD1 |
| Monocytes | CD14, FCGR3A, CD4 |
| Dendritic cells | FCER1A, CST3 |
| Melanocytes | PMEL, MLANA, TYR |
| Endothelial cells | CD93, CD34, LYVE1 |
| Monocytes | CD14, FCGR3A, FCGR3B |
| CAF | FAP, PDGFRA, PDGFRB, TAGLN, COL1A1 |
| alpha | GCG |
| beta | INS |
| delta | SST |
| gamma | PPY |
| epsilon | GHRL |
| ductal | KRT19 |
| acinar | CPA1 |
| endothelial | KDR, ESAM, FLT1, CDH5 |
| mesenchymal | SERPINE1 |

Markers for cell type identification in Seurat analysis

### Supplementary Table 2

| Datasets | Cell types |
| --- | --- |
| Baron | alpha, beta, delta, gamma, ductal, acinar, epsilon, endothelial, macrophage, schwann, mast, t cell, activated_stellate, quiescent_stellate |
| Muraro | alpha, beta, delta, gamma, ductal, acinar, endothelial, mesenchymal |
| Segerstolpe | alpha, beta, delta, ductal, acinar, mesenchymal, other |
| Wang | alpha, beta, gamma, delta, ductal, acinar, endothelial, mesenchymal, ambiguous |
| Xin | alpha, beta, gamma, delta, contaminated alpha, contaminated beta, contaminated gamma, contaminated delta |

List of cell types in pancreas benchmark

### Supplementary Table 3

| Datasets | Cell types |
| --- | --- |
| Sade-Feldman melanoma | NK, B cells, T cells, Monocytes, DC |
| PBMC 3k | NK, B cells, T cells, Monocytes, DC, Platelet |
| PBMC 500 | NK, B cells, T cells, Monocytes, DC |
| PBMC - Ding SM2 | NK, B cells, T cells, Monocytes, Megakaryocyte |
| HIV1 | NK, B cells, T cells, Monocytes, DC |
| Lung - Zilionis | NK, B cells, T cells, Monocytes, DC, Eosinophils, Neutrophils, Mast cells |

List of cell types in closely related populations  benchmark

### Supplementary Table 4

| Cell types | Markers |
|---|---|
| alpha (1) | GCG, TTR |
| beta (1) | DLK1, IAPP, INS |
| delta (1) | RBP4, SST |
| gamma (1) | PAX6, PPY |
| acinar (1) | CPA1, CPA2, CTRB2, PRSS1, SERPINA3 |
| ductal (1) | CFTR, KRT17, KRT19, SERPINA3 |
| B cells (1) | CD19, MS4A1, SDC1, CD79A, CD79B, CD38, CD37, CD83, CR2, MVK, MME, IL2RA, PTEN, POU2AF1, MEF2C, IRF8, TCF3, BACH2, MZB1, VPREB3, RASGRP2, CD86, CD84, LY86, CD74, SP140, BLK, FLI1, CD14, DERL3, LRMP |
| T cells (1) | ITGAL, CD4, CD44, TNFRSF9, GZMB, CD69, KLRB1, CCR6, CD2, CCR7, IL2RA, CD27, CD3G, CXCR5, ICOS, PLD4, CD3D, IL7R, CXCR6, CD28, CCR4, CCR10, CXCR3, SELL, CD3E, LTB |
| NK (1) | CD3E, CD3D, NCAM1, NCR1, GNLY, NKG7, IL32, CD27, FCGR3A, KLRD1, IL7R, PTPRC, TBX21 |
| Monocytes (1) | TYROBP, FCN1, FTL, TLR8, TLR4, ACE, CD14, PSAP, FCGR3A, PECAM1, ADGRE1 |
| Dendritic cells (1) | KLRD1, GSN, CD14, FCER1A, NCR1 |
| Dendritic cells (1*) | expressed: GSN, FCER1A<br>not expressed: KLRD1, CD14, NCR1 |
| B cells (2) | IGKC, MS4A1, IGHM, CD79A, CD19, CD22, BANK1, AC096579.7, BCL11A, FCRL1 |
| T cells (2) | CD8A, TRAC, CD8B, CD3D, CD3G, CD2, CTLA4, ICOS, IL32, ITM2A |
| NK (2) | TRDC, TRDV1, GNLY, FGFBP2, TRGC1, TRGC2, CTSW, GZMB, KLRD1, GZMH |

| | |
|---|---|
| Monocytes (2) | RP11-1143G9.4, LYZ, CST3, SERPINA1, FCER1G, TYROBP, CD14, AIF1, PLAUR, S100A9 |
| Dendritic cells (2) | LILRA4, SERPINF1, PLD4, IGJ, IL3RA, RP11-38J22.6, PTPRS, SPIB, TSPAN13, SMPD3 |

Markers used in training models and classifying cells

(1) for scClassifR and Garnett in pancreas benchmark, SCINA-1 in closely related cell types benchmark

(1*) for Garnett-1 in closely related cell types

(2) for SCINA-2 and Garnett-2 in closely related cell types
